# Cerebello-basal ganglia networks and behavior across adulthood and aging

**DOI:** 10.64898/2026.10.26.760492

**Authors:** Casey A. Delaney, Ivan A. Herrejon, Thamires N. Magalhaes, T. Bryan Jackson, Tracey H. Hicks, Jessica A. Bernard

## Abstract

The cerebellum contributes to aging, cognition, and reward related processing, but has been understudied in these domains. The cerebello-basal ganglia (CB-BG) circuit may be key to better understanding how the brain ages, as it has been implicated in modulation of cognitive, motor, and reward-related functions. Prior work has implicated this circuit in aging in a small sample, though it is also of interest in age-related neurodegenerative disease, particularly Parkinson’s Disease. Here, we investigated the CB-BG network in a sample of 138 participants between the ages of 35 and 86. Participants completed motor and cognitive testing batteries followed by 26 minutes of resting state functional connectivity magnetic resonance imaging (fcMRI), replicating previously mapped cognitive and motor subcircuits. When examining connectivity by age group (early and late middle age and older adulthood), we found higher functional connectivity in early middle age compared to older adulthood, with no results related to late middle age. Analysis of connectivity related to Stroop task performance showed higher connectivity-behavior associations in early middle-aged adults compared to older adults. Regarding sex differences, we found lower functional connectivity in females compared to males. Analysis of steroid hormone levels revealed higher connectivity associated with progesterone in females relative to males. We have replicated CB-BG subcircuits and further elucidated connectivity patterns associated with age, sex, progesterone levels, and cognitive performance. This expanded investigation of CB-BG networks represents an essential step in increasing our knowledge of age-related cognitive and motor declines and may be particularly relevant for consideration in Parkinson’s Disease.

## Introduction

With the rapidly aging population, it is critical to better understand the neural underpinnings of cognitive and motor age-related declines and identify new potential targets for remediation. While the cerebral cortex and hippocampus have long been the primary focus of aging research, they do not cover the full scope of changes to the aging brain and behavior. A growing literature has now implicated the cerebellum as an important area of investigation due to its role in cognitive and motor behaviors (Chen & Desmond, 2005b, 2005a; Doyon & Ungerleider, 2002; Imamizu et al., 2000; Leiner et al., 1986, 1989, 1991, 1993; Stoodley & Schmahmann, 2009) that are often impacted by age (Bernard et al., 2013a; Bernard & Seidler, 2013; Ma et al., 2025; McElroy et al., 2024; Seidler et al., 2010). Across healthy aging, the cerebellum undergoes structural changes in its total volume (Hoogendam et al., 2012; Jernigan et al., 2001; N. J. Lee et al., 2009; Walhovd et al., 2005, 2011), vermis volume (Raz et al., 1998, 2001; Rhyu et al., 1999; Shah et al., 1991; Sullivan et al., 2000), gray matter (Andersen et al., 2003), and white matter (Abe et al., 2008; Bennett et al., 2010; Giorgio et al., 2010). Deficits in cerebellar morphology have been associated with decreases in motor performance (Raz et al., 2000; Rosano et al., 2007; Woodruff-Pak et al., 2001) and decreased cognitive function (Hogan et al., 2011; J.-Y. Lee et al., 2005; MacLullich et al., 2004; Miller et al., 2013; Paul et al., 2009). Cerebellar functional changes are also prevalent, with changes observed in functional activation (Bo et al., 2011; Jackson et al., 2020; Mattay et al., 2002; Taniwaki et al., 2007; Ward & Frackowiak, 2003) as well as resting state functional connectivity (Andrews-Hanna et al., 2007; Ballard et al., 2023; Damoiseaux et al., 2008; Hicks et al., 2024, 2025; Langan et al., 2010; Magalhães et al., 2025; Wu et al., 2007). These structural and functional differences across age are also associated with alterations in cognitive and motor performance (Andrews-Hanna et al., 2007; Bernard et al., 2013b; Bernard & Seidler, 2013; Hausman et al., 2020; Langan et al., 2010), highlighting the need to further study the cerebellum in aging.

The basal ganglia, while investigated in the context of dopaminergic contributions to cognition (Braunlich & Seger, 2013; Grahn, 2009; Haber, 2014) and in the context of Parkinson’s Disease (PD) (Frank, 2005; Mattay et al., 2002; Owen et al., 1998), are relatively understudied in cognitively normal aging. Anatomical, physiological, and behavioral evidence has uncovered distinct neuronal loops of the basal ganglia that are associated with motor and cognitive functions (Arsalidou et al., 2013; Leisman et al., 2014; Leisman & Melillo, 2013; Middleton & Strick, 2000), yet only a few studies have begun assessing how these functions may be influenced by aging. The basal ganglia show similar aging patterns to the cerebellum, with deficits in structural connectivity (Radhakrishnan et al., 2023; Q. Wang et al., 2010), functional activation (Xiong et al., 2022), and functional connectivity (Bo et al., 2014; Griffanti et al., 2018; Rodriguez-Sabate et al., 2021). While studies are limited, research has shown that differences in the basal ganglia with age were associated with reduced cognitive performance through use of structural and functional MRI (Bo et al., 2014; Langenecker et al., 2007; Rodriguez-Sabate et al., 2021; Xiong et al., 2022), diffusion MRI (Radhakrishnan et al., 2023), and post mortem assessment of amyloid beta protein deposition and lacunes, which are cerebrospinal fluid filled infarcts (Gold et al., 2005). Regarding motor differences, one review assessed the potential influence of the aging basal ganglia on mobility (Clark et al., 2019). Using event related fMRI, one group found reductions in basal ganglia function for motor switching with age (Coxon et al., 2010). These differences are highly relevant as reductions in basal ganglia functional connectivity are seen in patients with Parkinson’s Disease (Kwak et al., 2010; Rodriguez-Sabate et al., 2019; Rolinski et al., 2015; Szewczyk-Krolikowski et al., 2014).

The cerebellum and basal ganglia both contribute to motor and cognitive behavior on their own and are structurally and functionally connected to each other in both primates and humans as a cerebello-basal ganglia (CBBG) circuit (Bostan et al., 2010, 2013; Bostan & Strick, 2010, 2018; Hausman et al., 2020; Milardi et al., 2016; Pelzer et al., 2013). Like each region alone, this CBBG circuitry is implicated in cognitive and motor behavior (Bernard et al., 2013a; Jackson & Bernard, 2022a, 2023; Middleton & Strick, 2000). Moreover, there is evidence that the structural connectivity between the cerebellum and basal ganglia is impacted by aging (Radhakrishnan et al., 2023). Previous work from our lab has shown, cross-sectionally, that there is lower CBBG FC in older adults than in young adults (Hausman et al., 2020) and CBBG FC is associated with cognitive and motor processing. Specifically, FC between Crus I and superior ventral striatum (VSs) was associated with Montreal Cognitive Assessment (MOCA) (Nasreddine et al., 2005) scores, and in older adults only, there was a relationship between these regions and self-reported balance confidence (Hausman et al., 2020). This was an essential first step in understanding the contributions of this circuit in healthy aging and its relationship to cognitive and motor processes, but further research is necessary to uncover the full influence of the CB-BG circuit.

Females represent an at-risk population for a decline in motor performance (e. g. falling) in advanced age (Peel et al., 2002; Stevens & Sogolow, 2005; Wei & Hester, 2014; Wolfson et al., 1994) and for Alzheimer’s Disease (AD) (Inzitari et al., 2006; Paganini-Hill & Henderson, 1994; Pike, 2017; Wray & Blaum, 2001), while males are at higher risk for developing Parkinson’s Disease (Cerri et al., 2019). There are reported sex differences in aging in the basal ganglia, though findings are not consistent across studies. Some groups report the right putamen and pallidum showing lower volumes in males than females (Y. Wang et al., 2019). Larger caudate gray matter volumes have been observed for males (Filipek et al., 1994), though these results disappear when controlling for brain size (Luders et al., 2009). In the cerebellum, it has been reported that men experience higher declines in gray matter volume in Lobules VIIIA, right lobules I-III, and Vermis IX and X but less declines in bilateral Lobules VIIIA (Han et al., 2020). Sex differences across aging may be potentially attributed to hormonal influences (Ballard et al., 2023; Hedges et al., 2012; Magalhães et al., 2025). Estrogen is highly relevant in neurotransmission, acting to influence physiological properties of a neuron and as a neurotransmitter (Hedges et al., 2012). Additionally, estrogen may be neuroprotective and influences female brain structure and circuits during stages of hormonal transitions, like post-partum or perimenopause (Chechko & Nehls, 2025; Russell et al., 2019). This neuroprotective effect has also been reported in risk of AD (Henderson & Brinton, 2010; Uddin et al., 2020). As the cerebellum contains dense receptors for estrogen (Almey et al., 2015; Hedges et al., 2012; Marrocco & McEwen, 2016), declines in estrogen levels and function across age may be a key factor influencing differences in cognitive and motor performance (Boyle et al., 2020; Sipilä et al., 2015). Further, this may impact CBBG connectivity, as we have previously demonstrated impacts on cerebello-cortical networks(Ballard et al., 2023; Hicks et al., 2025; Hicks & Bernard, 2025; Magalhães et al., 2025).

Here, we initially set to replicate previous work from our own group in an independent larger sample and further map resting-state functional connectivity between the cerebellum and the basal ganglia (Bernard et al., 2012; Hausman et al., 2020), investigating impacts of age and sex on these circuits. We also explored the impact of aging on FC in motor and cognitive behavior and respective sub-circuits that were defined using previous work assessing associations between functional connectivity and behaviors. We hypothesize that (i) increases in age will be associated with lower FC; (ii) FC patterns of cognitive and motor subcircuits will be correlated with their respective behaviors, and these subcircuits will be sensitive to aging; (iii) there will be higher FC in females compared to males; (iv) there will be an association between sex steroid hormones and FC in females compared to males in the context of aging. The present study will provide new insights into our understanding of the understudied CB-BG circuit, allowing us to assess FC changes over healthy aging, understand its relationship to cognitive and motor processes, and uncover regions at risk during pathological aging.

## Methods

### 1.1. Study sample

Participants (total *n*=175) were enrolled in a larger study on aging. During this assessment, all participants underwent a battery of cognitive and motor tasks, which are described in detail below. After the behavioral visit, the participants returned for a magnetic resonance imaging (MRI) session approximately two weeks later. However, due to unexpected delays related to the COVID-19 pandemic, the time between the two sessions varied between participants (average 39.0 days ± 21.4 days).

Exclusion criteria were history of neurological disease, stroke, or formal diagnosis of psychiatric illness (e.g., depression or anxiety), contraindications for the brain imaging environment, diagnosis of mild cognitive impairment or dementia, use of hormone therapy (HTh; inclusive of testosterone therapy) or hormonal contraceptives (hormonal intrauterine device (IUD), possible use of continuous birth control (oral)), and no history of hysterectomy in the past 10 years. These latter exclusions were made to evaluate the impacts of normative endocrine aging on healthy adult females. For our analysis, we focused only on those with available neuroimaging and behavioral (cognitive and motor) data. Thus, our final sample included 138 participants (74 females (M = 56.70 years, SD = 14.28) and 64 males (M = 56.94 years, SD = 12.52)) seen in Table 1. All study procedures were approved by the Institutional Review Board at Texas A&M University, and written informed consent was obtained from each participant prior to initiating any data collection.

**Table 1.**
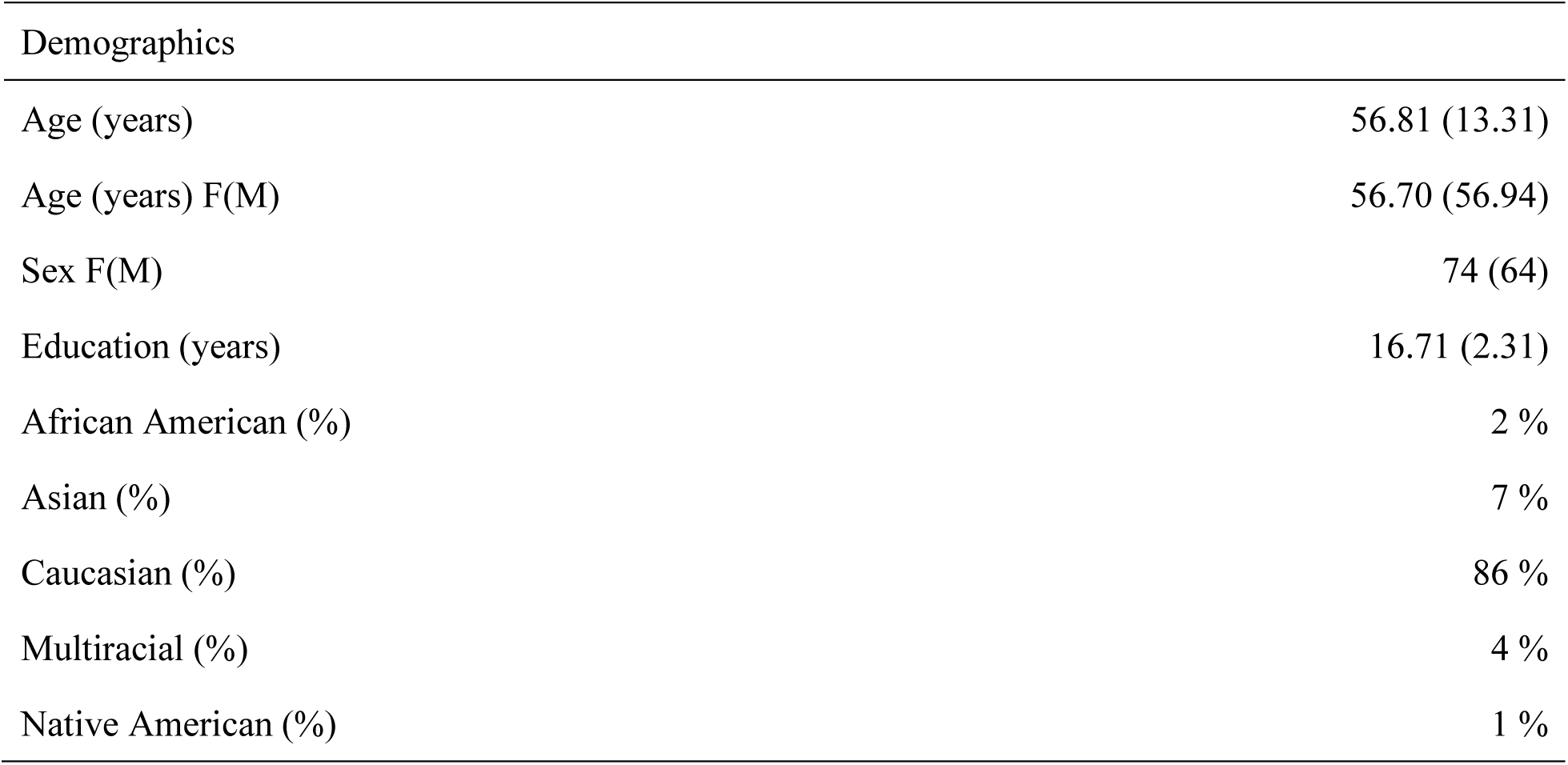
Demographic means for the full sample are listed below with standard deviations in parentheses.

### 1.2 Behavioral Protocol

#### 1.2.1 Cognitive Tasks

During the participants’ initial visit, they completed a battery of cognitive and motor tasks. They started by completing the Montreal Cognitive Assessment (MoCA) (Nasreddine et al., 2005). The Stroop task (Stroop, 1935) was completed via a computer using PsychoPy software. Participants were asked to identify the font color of a word rather than reading the word itself. Initially, participants completed a practice period where they received feedback on their accuracy. They then moved on to the testing phase, where they were given 1.5 seconds to respond to a stimulus on screen before the next stimulus appeared, without any time between trials. The response was categorized as a miss if a participant did not respond in time. Participants completed 10 blocks. Each block contained 18 congruent and 18 incongruent trials, with 360 total trials in the task. The Shopping List Memory Task (Flegal & Lustig, 2016) assessed longer term memory using a more ecologically realistic approach in asking participants to remember a shopping list. Participants were presented with one item at a time for three seconds and asked to memorize 15 items. After a 15-minute delay, participants were shown 15 items they had seen before and 15 new items. They were then asked to respond as to whether the item shown was an old or new one. Scores were calculated as the percentage of correct responses.

Participants also completed several subtests from the Wechsler Adult Intelligence Scale, 4th Edition (WAIS-IV): Digit Span, Letter-Number Sequencing, and Symbol Span. The Digit Span task consists of three conditions: participants read a string of numbers and recall the sequence in the same order as given, reverse order from given, or in ascending order. The letter-number sequencing test asks participants to listen to a sequence of numbers and letters and then recall the numbers in ascending order followed by the letters in alphabetical order. During symbol span, subjects are shown a series of symbols and then are asked to recall them in the order they were shown to them.

Based on previous work from our group (e.g., (Hicks et al., 2024)), two cognitive composite variables were created. The first composite included letter-number sequencing, the Stroop task, Symbol Span, and Digit Span to create a measure of working memory, executive functioning, processing speed, and attention. The second composite variable included the sequence learning task and the Shopping List Memory Task as a measure of episodic memory and motor learning. To create these composite variables, raw scores from each task were z-scored and then averaged together for each subject (Table 2). This approach allowed us to investigate associations between behavior and connectivity across domains while limiting multiple comparisons.

**Table 2.**
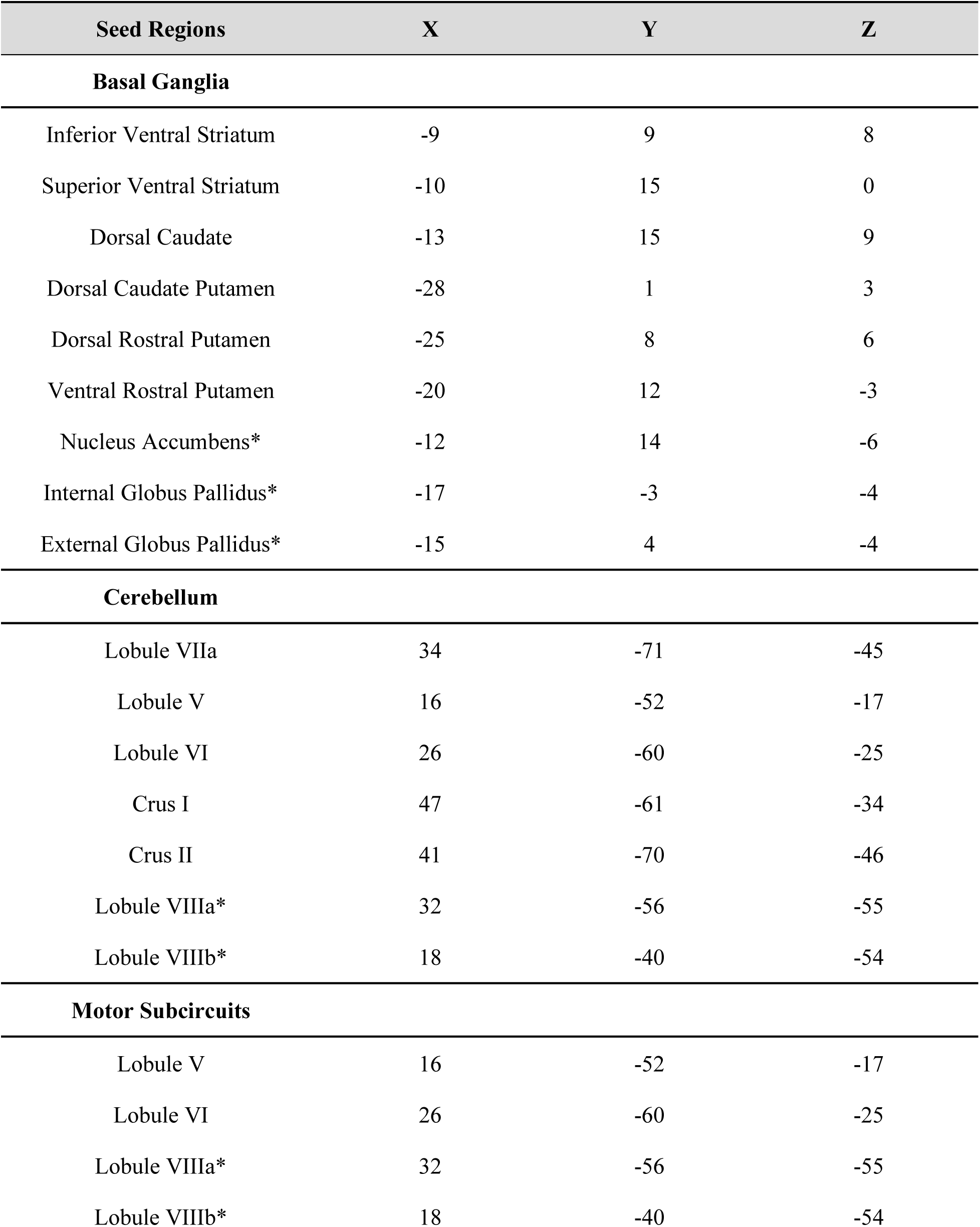
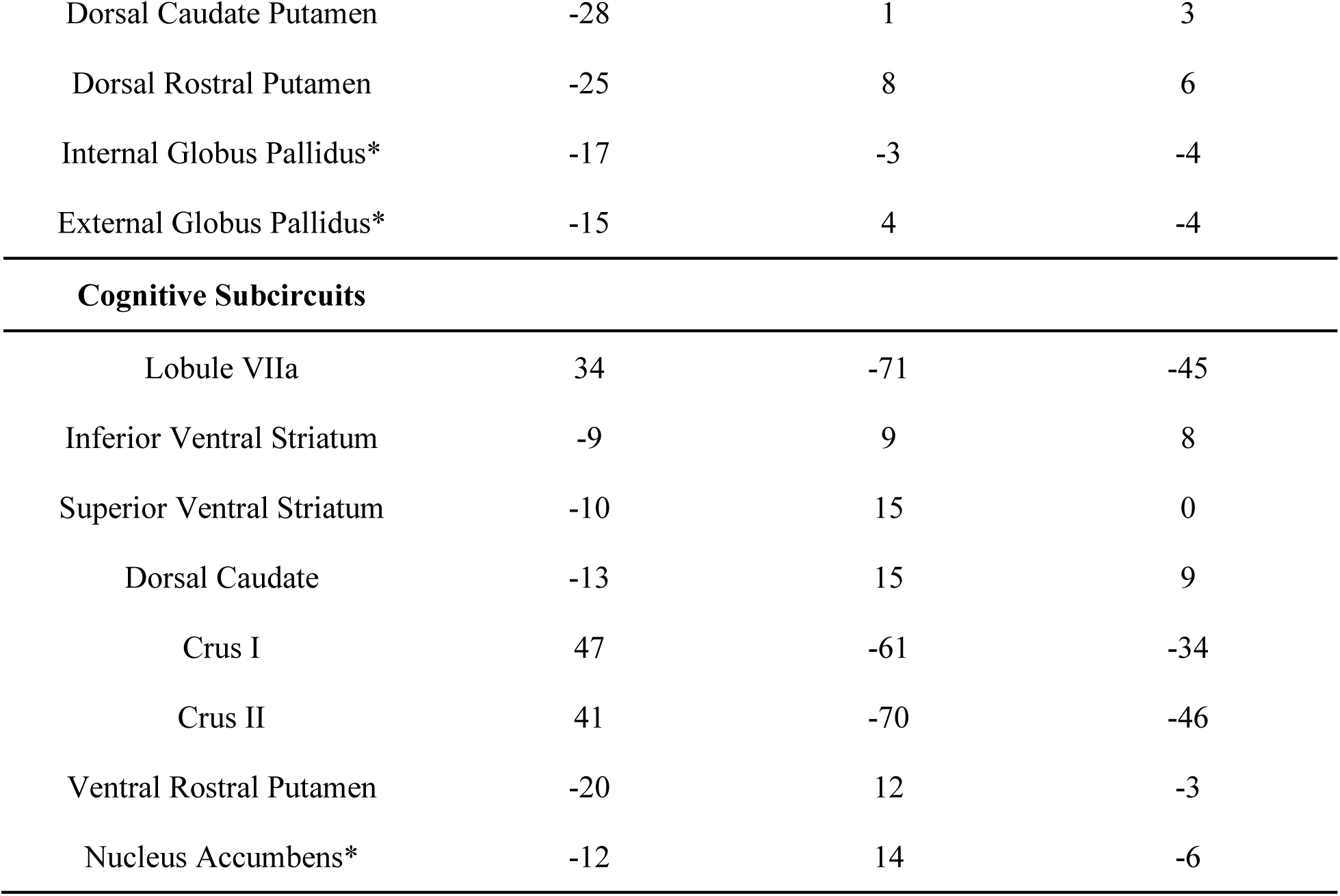
Coordinates for the cerebellum and basal Ganglia seeds organized by region and subcircuit. These were retrieved from Hausman, 2020. The ones denoted with a * symbol were new seeds added for these analyses to expand our understanding of this circuit.

#### 1.2.2 Motor Tasks

All participants completed the Purdue Pegboard Task (Tiffin & Asher, 1948), which consists of four conditions. Participants were given a set of pegs, tubes, and washers and asked to fill the pegboard. In the first condition, participants filled the left section of the pegboard using their left hand for 30 seconds. In the second, participants filled the right section with their right hand only for another 30 seconds. In the third condition, participants filled both the left and the right side with their left and right hand, respectively for 30 seconds as well. Finally, participants created an assembly using all four materials given, for 60 seconds. These four conditions were repeated three times. For the first three conditions, the number of pins placed correctly were counted. For the assembly condition, each component of the assembly was counted even if the assembly was not finished with all four parts. The first three conditions measure fine motor skills, while the last condition is used to test sequencing skills, bimanual coordination, and fine motor skills.

During the sequence learning task, participants were shown four squares on a computer screen that were shaded black, one at a time. Participants were asked to indicate which square was shaded by pressing a key on a keyboard. Participants were either shown a sequence that repeated itself or the order in which the blocks were shaded was random. Learning was quantified by calculating the slope of task accuracy from Block 1 to 9 of the sequence condition. The approach for calculating the learning slope was detailed in our recent work (Hicks et al., 2024).

Postural sway was quantified using an Advanced Mechanical Technology Incorporated Accusway force platform (Watertown, MA). Participants were asked to complete four conditions by staying as still as possible with their hands crossed across their chests. They were asked to stand with their eyes open and feet apart (EOOB), eyes closed and feet apart (ECOB), eyes open and feet together (EOCB), and lastly, eyes closed and feet together (ECCB). For two minutes with a sample rate of 50 Hz, the center of pressure (COP) was recorded. We applied a 9th-order Butterworth filter with a 20 Hz cutoff frequency to isolate the low-frequency postural sway. The 95% confidence interval of COP area was calculated consistent with previous literature (Oliveira et al., 1996; Sansare et al., 2024).

Finally, we conducted a bilateral grip strength test to measure gross motor upper limb function using a handheld dynamometer. Here, participants were asked to squeeze the handle as hard as possible with their dominant hand for five seconds to measure the kilograms of force of a grip.

#### 1.2.3 Hormone Collection

Saliva samples were collected on the day of the first visit for hormonal testing (Ballard et al., 2023; Hicks et al., 2025; Hicks & Bernard, 2025; Magalhães et al., 2025). We followed the recommended protocol from SALIMETRICS. Prior to saliva collection, participants were asked to fast for at least 60 minutes, to document the consumption of alcohol, nicotine, coffee, and/or medication 12 hours before collection, to indicate any oral injury or disease, and to rinse their mouth to remove possible food residue 10 minutes prior to sample collection. Participants were then asked to provide 1 mL of saliva using the passive drool technique. After completion, the time and date were recorded, and the saliva sample was immediately frozen in a bio-freezer for stabilization at -80 degrees Fahrenheit. Salimetrics completed the assays to measure 17β-estradiol, progesterone, and testosterone levels.

### 1.3. Imaging acquisition

As noted, these data and analyses are part of a larger study. For the sake of clarity and consistency in reporting, our boilerplate methods here are identical to those reported in Hicks (2024). Participants underwent structural and resting-state MRI using a Siemens Magnetom Verio 3.0 Tesla scanner and a 32-channel head coil. For structural MRI, we collected a high-resolution T1-weighted 3D magnetization prepared rapid gradient multi-echo (MPRAGE) scan (repetition time (TR) = 2400 ms; acquisition time = 7 minutes; voxel size = 0.8 mm 3) and a high-resolution T2-weighted scan (TR = 3200 ms; acquisition time = 5.5 minutes; voxel size = 0.8 mm 3), each with a multiband acceleration factor of 2. For the resting-state imaging, we administered four blood-oxygen-level-dependent (BOLD) functional connectivity (fcMRI) scans with the following parameters: multiband factor of 8, 488 volumes, TR of 720 ms, and 2.5 mm^3^ voxels. Each fcMRI scan was 6 minutes in length for a total of 24 minutes of resting-state imaging, and scans were acquired with alternating phase encoding directions (i.e., two anterior to posterior scans and two posteriors to anterior scans). During the fcMRI scans, participants were asked to lie still with their eyes open while fixating on a central cross. In total, the acquisition of images takes about 45 minutes, including a 1.5-minute localizer. Scanning protocols were adapted from the multiband sequences developed by the Human Connectome Project (HCP) (Harms et al., 2018) and the Center for Magnetic Resonance Research at the University of Minnesota to facilitate future data sharing and reproducibility.

#### 1.3.1. Imaging processing

Much like the data collection described above, we use identical processing across all our data, and for consistency and transparency across our work, we have again used boilerplate text as initially described by Hicks (2024). The images underwent several preprocessing steps to prepare them for further analysis. Initially, they were converted from DICOM to NIFTI format and organized following the Brain Imaging Data Structure (BIDS, version 1.6.0) using the bidskit docker container (version 2021.6.14, https://github.com/jmtyszka/bidskit). Afterward, a single volume was extracted from two oppositely coded BOLD images to estimate B0 field maps using the split tool from the FMRIB Software Library (FSL) package (Jenkinson et al., 2012). Subsequently, the anatomical and functional images were preprocessed using fMRIPrep (version 20.2.3; for detailed methods, see https://fmriprep.org/), which includes automated procedures to align the functional volume with the anatomical image, correct for motion, correct field map distortions, segment the anatomical image into distinct tissues (e.g., gray matter, white matter, cerebrospinal fluid), remove the skull from the anatomical image, normalize the data to a common space, align motion-corrected functional volumes with the normalized anatomical image, and apply spatial smoothing.

##### 1.3.1.2. Functional analysis

As reported by Hicks and colleagues (2024), following preprocessing with fMRIPrep, the subsequent analyses were conducted using the CONN toolbox (version 21a) (Whitfield-Gabrieli & Nieto-Castanon, 2012). This involved additional processing to eliminate noise and artifacts and enhance data quality. Denoising in CONN typically comprises several stages, such as removing motion signals and regressing out confounding signals (e.g., signals from ventricles, white matter, and global signals). Motion information from fMRIPrep was utilized in CONN for this purpose. A 0.008-0.099 Hz bandpass filter was applied to eliminate high-frequency noise. The denoising step is crucial for enhancing the quality of functional connectivity data by minimizing artifacts and enhancing the ability to detect genuine functional connectivity patterns in the brain.

In the first level analysis, we included cerebellar and basal ganglia spherical seeds with a radius of 3.5 mm that were used in our previous work (Hausman et al., 2020). For the basal ganglia, these include the inferior and superior ventral striatum (VSi & VSs), the dorsal caudate (DC), dorsal caudal putamen (DCP), dorsal rostral putamen (DRP), and ventral rostral putamen (VRP). For the cerebellum, we included seeds in lobules V and VI, as well as Crus I and II. Additionally, we added cerebellar seeds in lobules VIIIa and VIIIb to investigate the secondary motor representation and in lobule VIIa to better represent the non-motor/cognitive circuit. For the basal ganglia, additional seeds were created in the internal and external parts of the globus pallidus, as well as the nucleus accumbens based on resting state functional connectivity and structural MRI studies (Günther et al., 2018; Tarcijonas et al., 2020). Cerebellar and basal ganglia seed coordinates are listed in Table 2. To define motor and cognitive sub-circuits, seeds were divided as follows based on our previous work (Hausman et al., 2020). Subcircuits, in Table 3, were defined by previous white matter mapping and resting state parcellation studies, as well as functional connectivity studies that analyzed each subregion of the cerebellum and basal ganglia in relation to motor and cognitive behaviors (Balsters et al., 2014; Di Martino et al., 2008; Draganski & May, 2008; Hausman et al., 2020; King et al., 2019; Salmi et al., 2010; Stoodley & Schmahmann, 2009, 2009). ROI-to-ROI analyses were conducted. Here, we explored (i) FC differences associated with age (EA>OA, MA>OA, & EA>MA); (ii) CBBG Motor and Cognitive subcircuit FC and their associations with behaviors (iii) FC differences associated with sex (Females>Males) (iv) FC of each individual age groups (EA, MA, OA) (iv) Associations with cognitive (Comp1, Comp2) and motor tasks (sequencing, pegboard, and balance performance) in all subjects and sub-circuits; (v) Finally, we explored any FC associations with hormones (progesterone, estradiol, and testosterone). All analysis was completed using parametric statistics. We analyzed our results using an analysis level false discovery rate correction where P < 0.05, considering all the pairwise comparisons in the analysis.

**Table 3.**
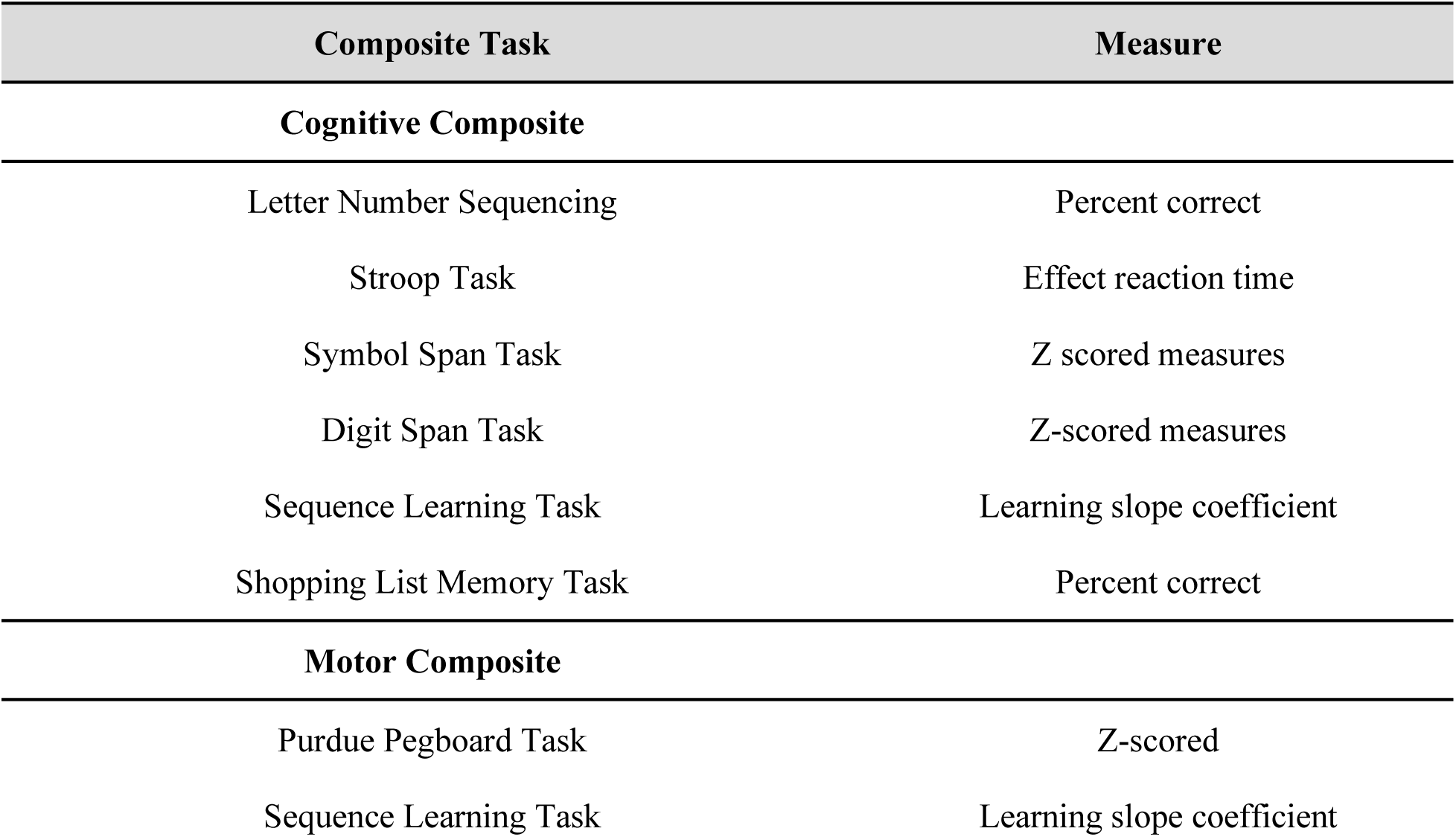
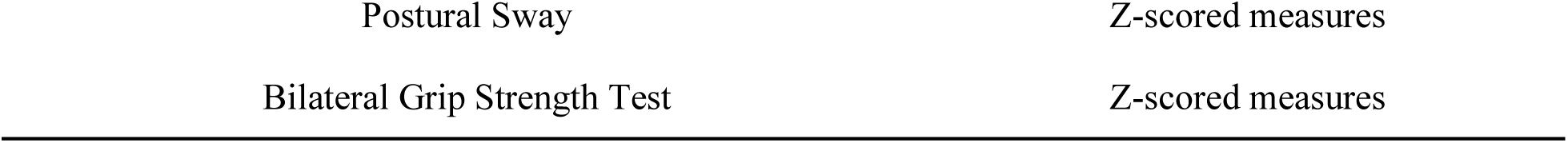
Cognitive and Motor Composites Tasks and Measures.

## Results

### The CBBG circuit can be elucidated into motor and cognitive subcircuits

Consistent with our previous work, we found that patterns of connectivity could be parsed based on cognitive and motor associations with the cerebral cortex, forming cognitive and motor subcircuits (Fig 1 & 2). These subcircuits varied in regions and strengths of connectivity between age groups (Fig 3 & 4). Areas of the cerebellum involved in cognitive processing and networks (Crus I & II) were highly correlated with areas of the basal ganglia involved in cognitive processing and networks (VSi, VSs, DC, VRP, and the nucleus accumbens). Similarly, areas of the cerebellum involved in motor processing (lobule V & VI) were highly correlated with areas of the basal ganglia involved in motor processing (DCP, DRP, GPi, and GPe).

**Figure 1.**
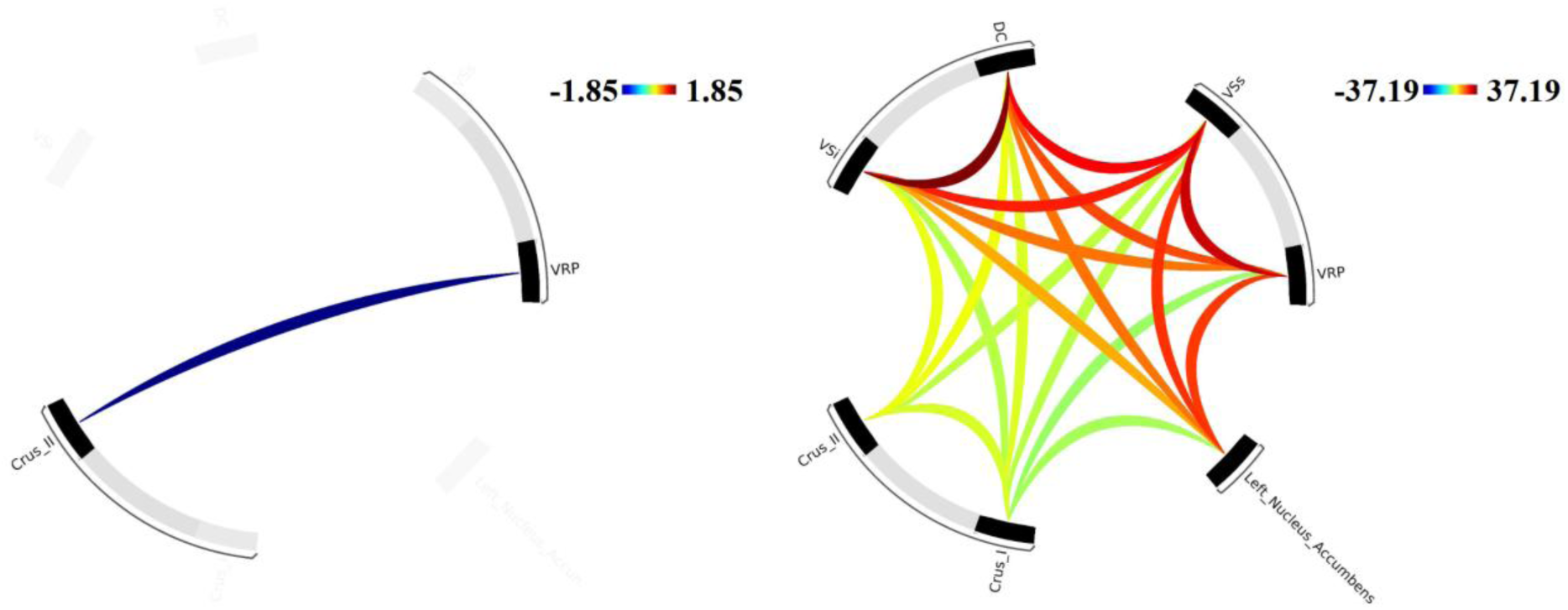
Patterns of functional connectivity (FC) between ROIs associated with cognitive behavior and networks. The color bar represents the strength of the connectivity with red being positive and blue being negative. Results were evaluated using an analysis level FDR correction such that P < 0.05.

**Figure 2.**
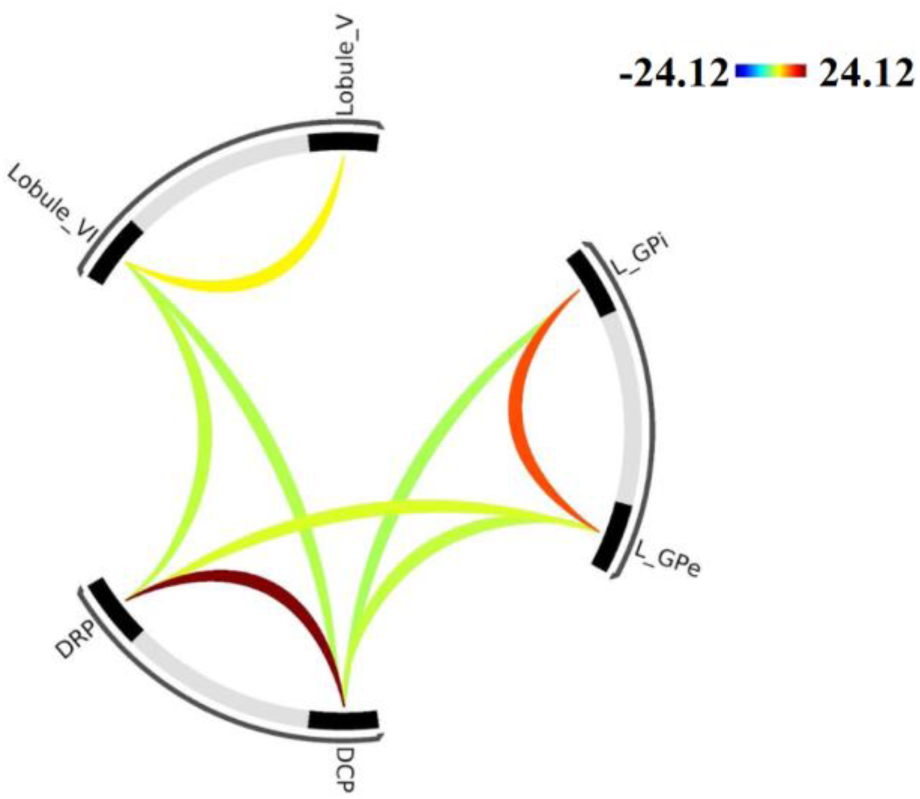
Patterns of functional connectivity (FC) between ROIs associated with motor behavior and networks. The color bar represents the strength of the connectivity with red being positive and blue being negative. Results were evaluated using an analysis level FDR correction such that P < 0.05.

**Figure 3.**
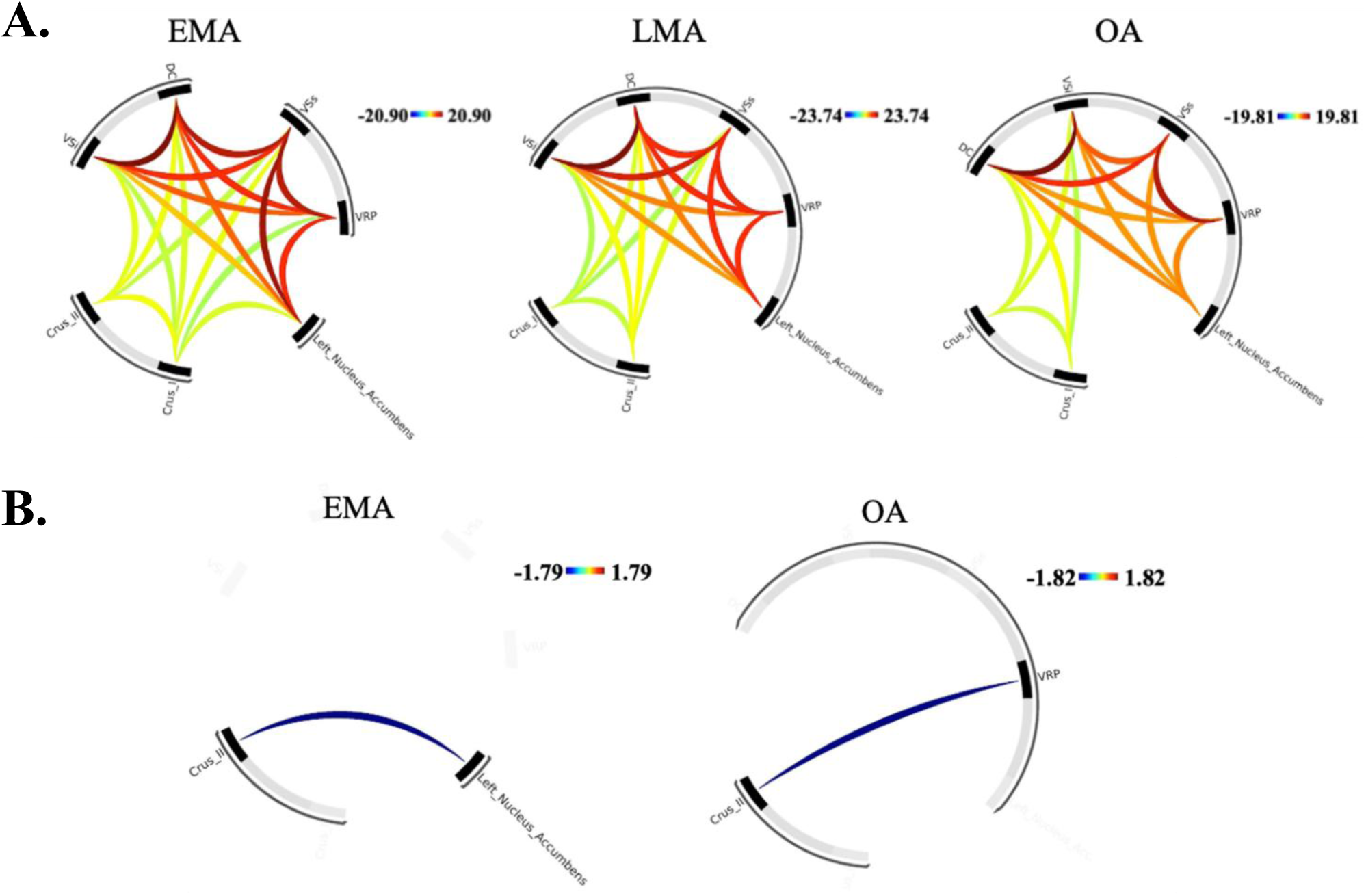
Patterns of functional connectivity (FC) of the cognitive subcircuit for each individual age group, including early middle age (EMA), late middle age (LMA), and older adulthood (OA). **A.** Positive connectivity. **B.** Negative connectivity. The color bar represents the strength of the connectivity with red being positive and blue being negative. Results were evaluated using an analysis level FDR correction such that P < 0.05. No lower connectivity between ROIs was seen for LMA.

**Figure 4.**
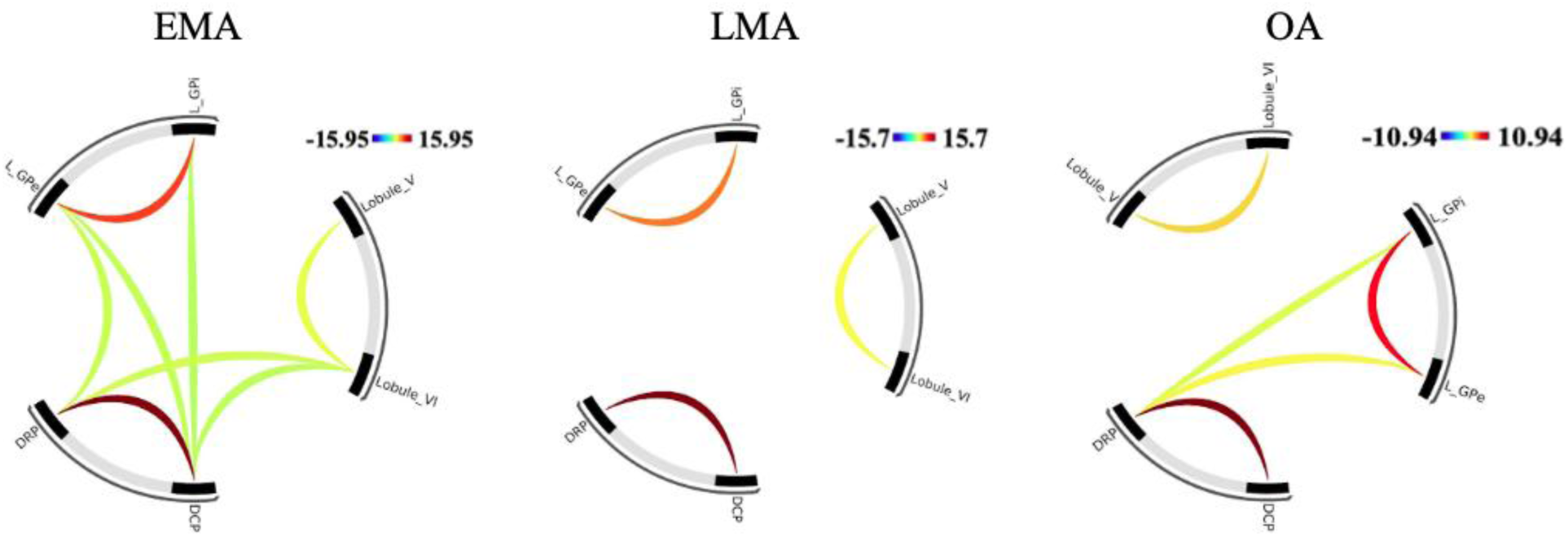
Patterns of functional connectivity (FC) of the motor subcircuit for each individual age group, including early middle age (EMA), late middle age (LMA), and older adulthood (OA). The color bar represents the strength of the connectivity with red being positive and blue being negative. Results were evaluated using an analysis level FDR correction such that P < 0.05. No negative associations between ROIs were seen in any age group.

### Functional connectivity of the CBBG circuit decreases with age

When contrasting OA>EMA, we found that connectivity of the CBBG circuit of the older adult group was significantly altered relative to early middle-aged adults (Fig. 5). Seen in Figure 2, we saw larger patterns of connectivity for Left Nucleus Accumbens of the basal ganglia with Crus I of the cerebellum. Additionally, the Lateral GPe was highly connected with the VSs, VSi, and DC. We also saw a high connectivity between the DC and the Lateral Gpi. Notably, we saw some regions of connectivity that were significantly higher for older adults when contrasted to early middle-aged adults. We saw that Lobule VIIIa was highly correlated with the VSi, VRP, DRP, and DCP of the basal ganglia. There also was a strong connection between VSs and the Lateral GPi. We did not find any differences when statistically comparing middle aged to early middle aged, or when comparing older adults to middle aged, suggesting that large changes in connectivity occur gradually over longer periods of aging, rather than as robust changes throughout each stage aging.

**Figure 5.**
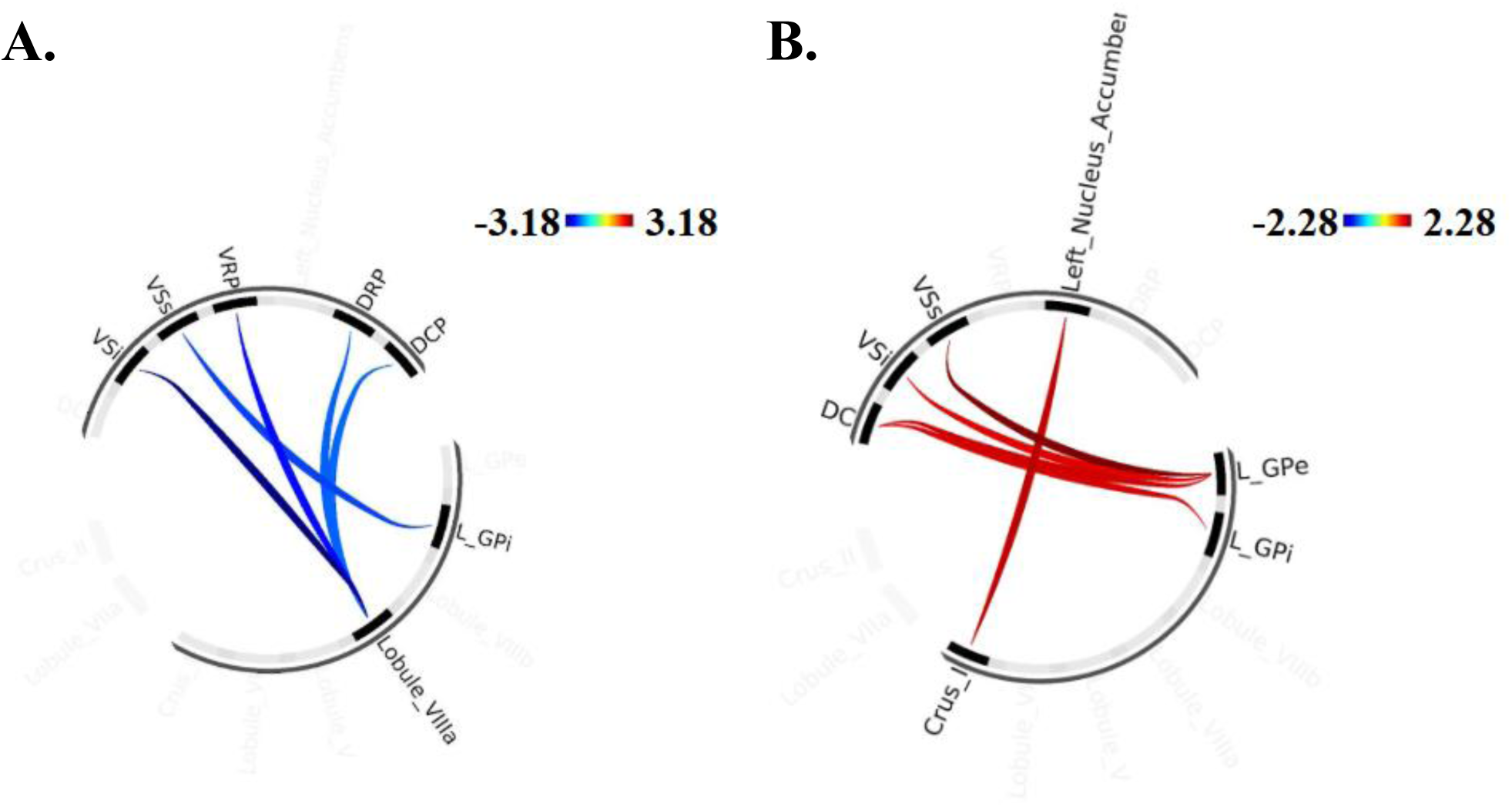
Age differences in cerebello-basal ganglia functional connectivity. **A.** Patterns of higher FC in the CBBG circuit for EMA relative to OA. **B.** Patterns of lower FC in the CBBG circuit for EMA relative to OA. The color bar represents the strength of the connectivity with red being positive and blue being negative. Results were evaluated using an analysis level FDR correction such that P < 0.05.

### Cerebello-basal ganglia connectivity is associated with cognitive behavior but not motor behavior

We found that CBBG connectivity was associated only with reaction times on the Stroop task (Fig 6), but not with the cognitive composite scores. Connectivity associations with behavior between Lobule VI and Crus II were lower in older adults relative to early middle-aged adults. We found higher connectivity between Lobule VIIIa with Crus I and the Lateral GPe, as well as higher connectivity of Crus I with Lobule VI and Lobule V in older adults relative to early middle-aged adults. We did not find associations of the CBBG cognitive subcircuit with the cognitive composite. All results were across all ROIs. We also did not find any associations of the CBBG connectivity with motor behavior.

**Figure 6.**
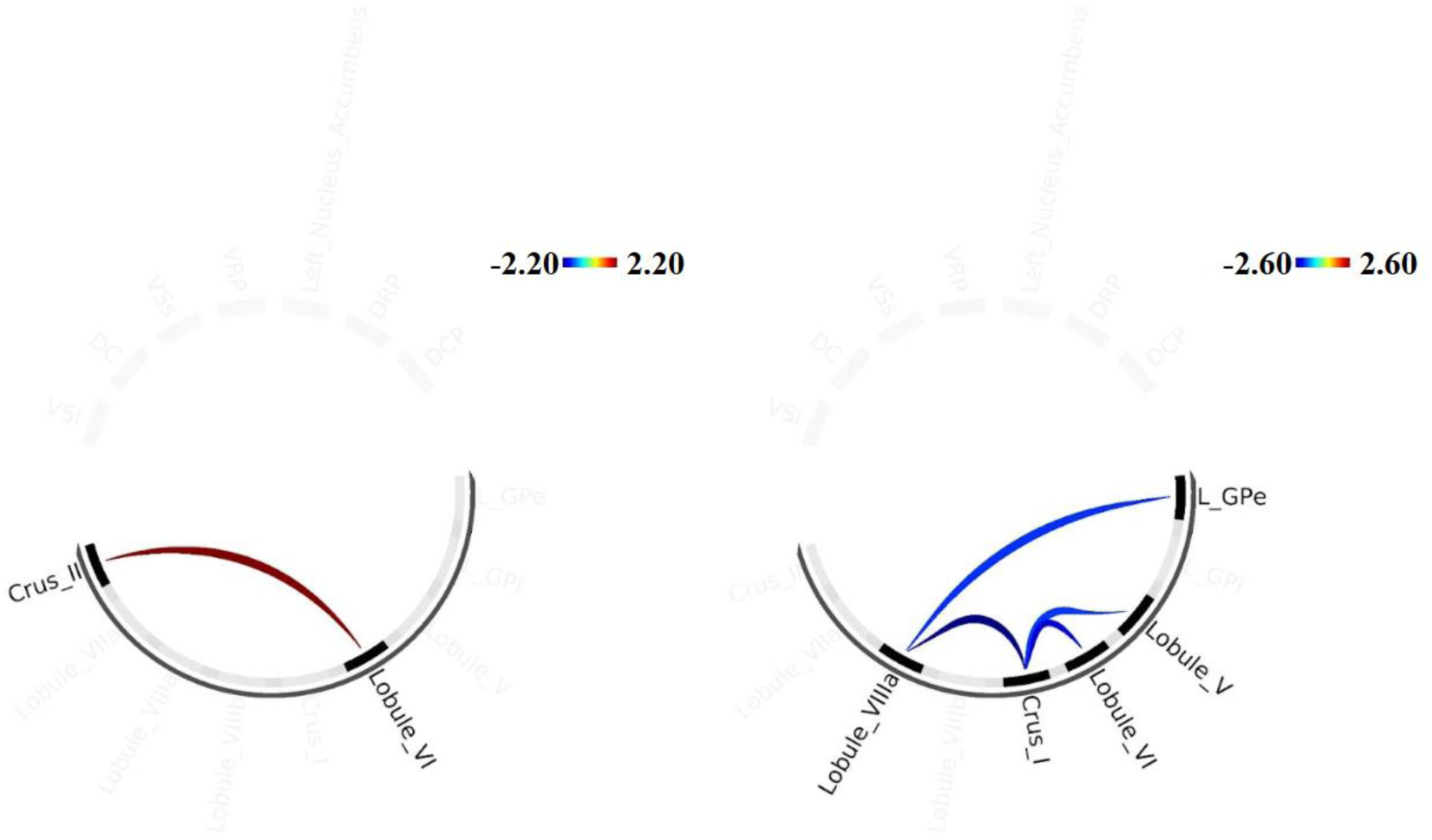
Differences cerebello-basal ganglia functional connectivity associations with cognitive behavior across age. **A.** Higher FC in the CBBG circuit associated with reaction times on a Stroop task for EMA relative to OA. **B.** Patterns of lower FC in the CBBG circuit associated with reaction times on a Stroop task for EMA relative to OA. The color bar represents the strength of the connectivity with red being positive and blue being negative. Results were evaluated using an analysis level FDR correction such that P < 0.05.

### Sex differences in CBBG connectivity and associations with progesterone

We found lower FC only *within* basal ganglia regions for females relative to males (Fig. 7), highlighting an interesting sex difference in circuitry across our sample. Further, we also found that CBBG connectivity was associated with progesterone differentially based on sex, suggesting this circuit is sensitive to hormones uniquely for males and females (Fig. 8). When contrasting Females>Males, males showed higher levels of connectivity *within* the cerebellum and *within* the basal ganglia associated with progesterone levels, and females showed higher levels of connectivity *between* the cerebellum and basal ganglia associated with progesterone levels. There were no significant associations with 17β-estradiol or testosterone.

**Figure 7.**
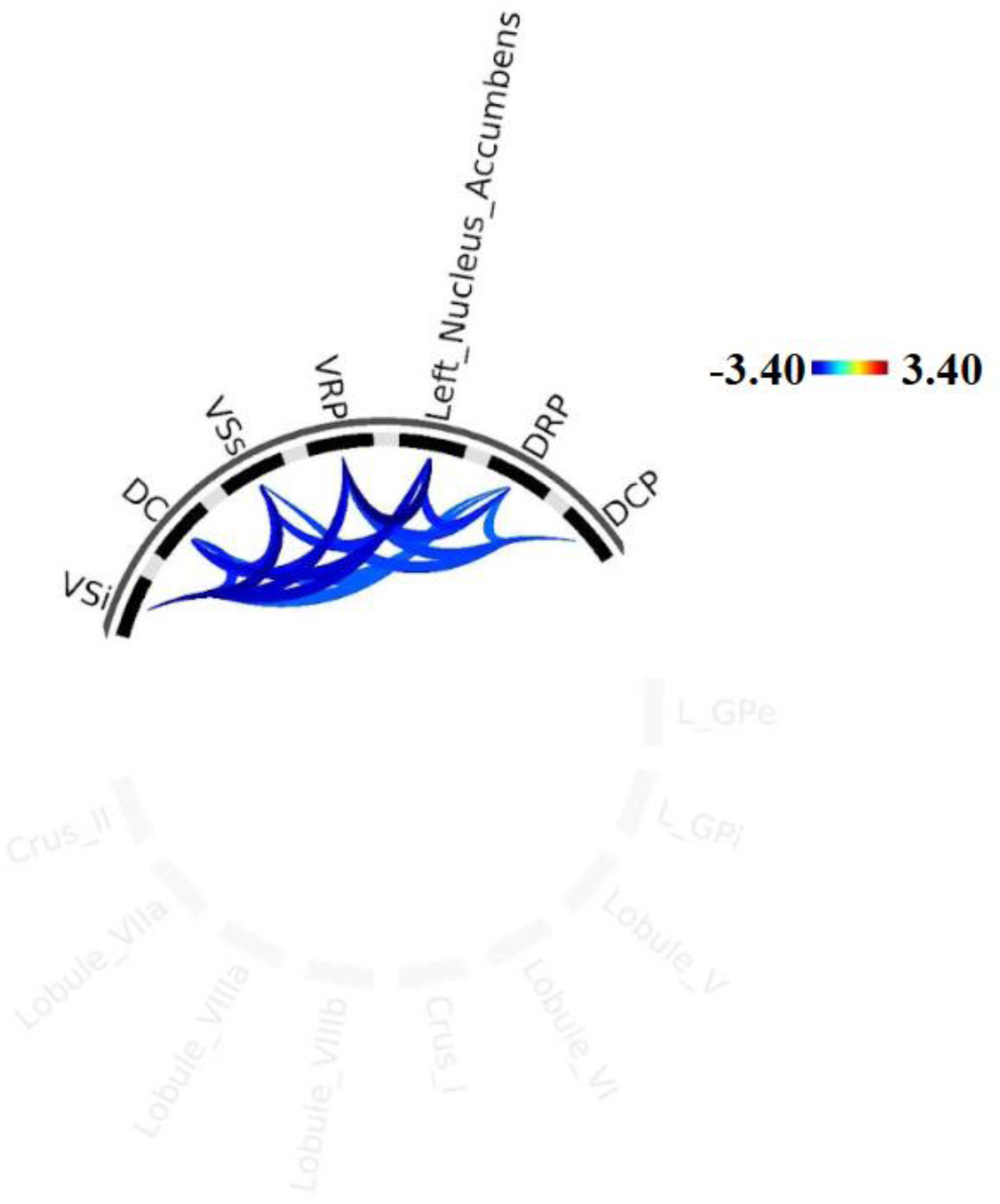
Sex differences in cerebello-basal ganglia functional connectivity. Patterns of lower FC in the CBBG circuit for females relative to males, that were localized within the basal ganglia. The color bar represents the strength of the connectivity with red being positive and blue being negative. Results were evaluated using an analysis level FDR correction such that P < 0.05.

**Figure 8.**
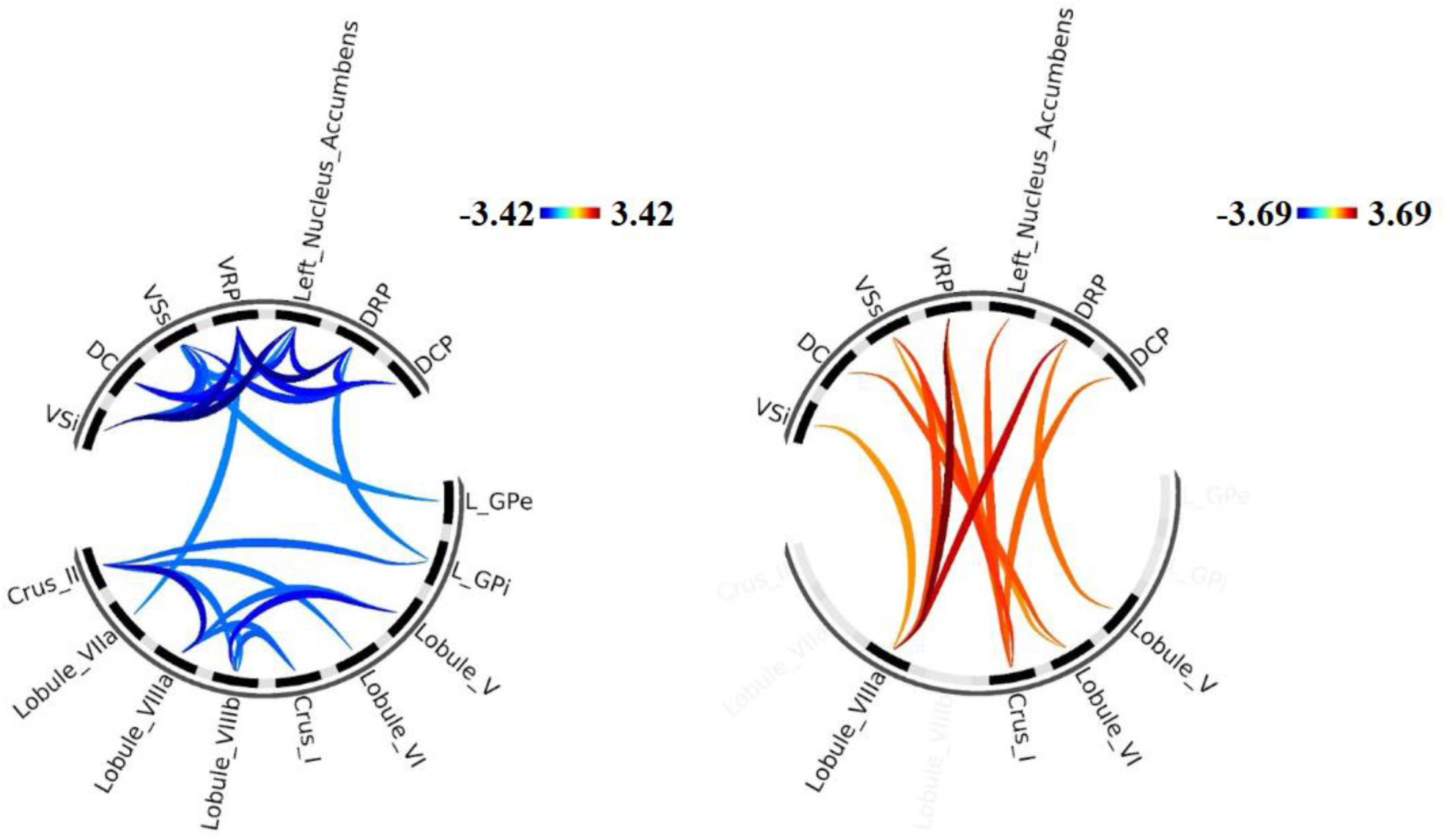
Differences in cerebello-basal ganglia functional connectivity associations with progesterone across sex, comparing Females>Males. The blue image represents areas of higher connectivity in males while the red image represents areas of high connectivity in females. The color bar represents the strength of the connectivity with red being positive and blue being negative. Results were evaluated using an analysis level FDR correction such that P < 0.05.

## Discussion

The cerebellum and basal ganglia are individually relevant in the context of cognitive and motor performance, especially across age (Andrews-Hanna et al., 2007; Arsalidou et al., 2013; Bernard et al., 2013b; Hausman et al., 2020; Langan et al., 2010; Leisman et al., 2014; Leisman & Melillo, 2013; Middleton & Strick, 2000). These regions are also functionally linked in a CBBG circuit(Bernard et al., 2013b; Hausman et al., 2020; Jackson et al., 2022; Jackson & Bernard, 2022a, 2022b, 2023; Middleton & Strick, 2000), highlighting a potential target of study for age-related motor and cognitive decline. No previous study has evaluated this circuit in the context of age, sex, and behavior. Using an ROI-ROI approach, we were able to replicate the circuit, it’s subcircuits, and evaluate how its functional connectivity is associated with motor and cognitive behavior, with increasing age across adulthood, sex, and sex steroid hormone levels (17β-estradiol, progesterone, and testosterone).

In line with our previous work (Hausman et al., 2020), we found that the CBBG circuit could be parsed into cognitive and motor subcircuits. We replicated our findings such that Crus I and Crus II of the cerebellum are functionally connected to VSs and DC of the basal ganglia in a cognitive subcircuit. Here, we also uncovered additional regions of the basal ganglia (VSi, DC, and VRP) that are functionally connected to the cerebellum. We further replicated our previous motor subcircuit, finding functional connectivity between Lobules V and VI of the cerebellum and the DRP and DCP of the basal ganglia. This functional connectivity further extended to the additional basal ganglia regions we added here, the GPi and GPe. These findings highlight the heterogeneous functional contributions of these regions. We have also expanded our understanding of key regions involved in these circuits, allowing for more precise assessment of specialized neural differences with age.

Previously, we assessed differences in CBBG functional connectivity only between young and older adults (Hausman et al., 2020). Here, we have included middle aged adults to understand the period of adulthood during which differences in brain connectivity may begin to emerge. We found that CBBG functional connectivity was lower in older adults when compared to early middle-aged adults. No significant differences in functional connectivity were seen when comparing early middle aged to late middle-aged adults or when comparing late middle aged to older adults. These findings suggest that differences in the CBBG circuit occur over time, with small differences across midlife that eventually amass to a larger difference in older age.

When evaluating differences in cognitive performance associated with FC and age, the only significant finding was reaction times on a Stroop task associated with FC. The comparison of older adults relative to early middle-aged adults also revealed differences in FC associations with Stroop task reaction time within the cerebellum only. This association was seen when investigating the full circuit, but not cognitive subcircuits and was also not seen for the whole cognitive composite score. This single CBBG-behavior association with reaction time may be related to the involvement of the cerebellum in the automaticity of tasks (Balsters & Ramnani, 2011; Thach, 1998) and inhibition of interfering information (Bellebaum & Daum, 2007). The CBBG circuit may be recruited when individuals require more effort to make a task automatic. Prior work showed that older adults were able to become automatic at a sequential motor task, though it took longer and required increased activation of the cerebellum compared to younger adults (Wu & Hallett, 2005). Another study assessed whole brain connectivity with cerebral blood flow during the Stroop task, finding that the cerebellum was significantly coactivated with the dorsal and ventral prefrontal cortex during incongruent trials (Harrison et al., 2005). They attribute this finding to the possibility that this task does not become less effortful, so the cerebellum was still activated. As Stroop accuracy did not show any associations with connectivity or age, it is possible that older individuals are able to continue performing at a high level, but at the cost of their time and effort, resulting in the correlation between CBBG connectivity and reaction time.

The cerebellum and basal ganglia have been hypothesized to operate as an internal timing system (Diedrichsen et al., 2003; Ivry & Keele, 1989). When assessing the involvement of these regions in prediction error and timing, one group found that the posterior cerebellum was involved in timing adjustment (Dreher & Grafman, 2002). Other studies have shown the involvement of the cerebellum and basal ganglia in visual movement timing in non-human primates, revealing that the cerebellum may be involved in precise regulation self-timing on a subsecond level, whereas the basal ganglia keep track of actual vs predicted time elapsed on a suprasecond scale (Kunimatsu et al., 2018; Tanaka et al., 2024). It is possible that regions of the cerebellum are engaged to maintain the self-timing of the motor response. As the basal ganglia are more involved in keeping track of time, they may be less recruited for a Stroop task, where timing of trials is consistent and does not need to be tracked.

Regions of higher connectivity associated with cognition in older adults may be an effect of compensation. Compensation suggests that older adults recruit additional neural resources that are not recruited in younger adults to compensate for a lack of connectivity elsewhere and maintain cognitive performance (Cabeza et al., 2018). The cerebellum has been posited as a form of reserve, activating as additional support in cognitive domains when prefrontal resources are exhausted(Bernard, 2022; Guo et al., 2016; Mitoma et al., 2020, 2022; Mitoma & Manto, 2023; Schmahmann, 2016; Tang et al., 2021). On a cortical level, it is possible that older adults recruit more of the CBBG circuit as compensation for a potential lack of prefrontal engagement. Subcortical compensation, such that different subsections of the CBBG circuit are recruited for older adults if the previously used regions were not sufficient to maintain performance, may be occurring. This is especially unique as compensation has mainly been studied in a cortical context (Mattay et al., 2002; Reuter-Lorenz & Cappell, 2008; Reuter-lorenz & Mikels, 2006; Tang et al., 2021), but our findings suggest that this may be happening subcortically as well. However, longitudinal investigations would provide a more robust test of this assertion, and here we were limited to cross-sectional data.

When investigating sex differences in CBBG circuitry, we saw lower FC only in basal ganglia regions for females. This finding is particularly surprising as reductions in basal ganglia connectivity have been associated with diagnosis of Parkinson’s Disease (Szewczyk-Krolikowski et al., 2014); PD), yet PD is typically more commonly diagnosed in males (Cereda et al., 2016). As the individuals in this study were healthy adults, it is possible that this pattern may be unique to healthy aging rather than pathological aging. Additionally, there are limited and often contradictory findings on sex differences in the basal ganglia across adulthood, with some groups reporting larger caudate gray matter volumes in males (Filipek et al., 1994), while others report the opposite after controlling for brain size (Luders et al., 2009). Another study reported reductions in the right putamen and right pallidum for aging males (Y. Wang et al., 2019). These studies show mixed results and only assess structural differences in the basal ganglia. Functional connectivity results are limited, though one group found a sex by age-squared interaction in the left caudate and strong correlation in subcortical network connectivity with age for females (Scheinost et al., 2014). Further work is needed to better understand how the connectivity of the basal ganglia is influenced by age and sex.

In males, there were positive associations between progesterone with FC within the basal ganglia and cerebellum and only limited interconnectivity between the cerebellum and basal ganglia. No associations were seen with estradiol, likely as there are not drastic fluctuations in this hormone in aging males. In females, we saw robust associations between progesterone levels and basal ganglia-cerebellar FC. This finding may be attributed to the menopause related drop in progesterone for females, such that there may have been greater progesterone variability in our female sample at assessment. Variability in the levels of estradiol and testosterone in the female sample at assessment may also explain why no associations were found with either hormone. Testosterone also does not fluctuate as drastically as other hormones in aging females, so there may be less influence on brain connectivity (Davis et al., 2019). Further, this study did not exclude females on any hormone replacement therapy, so it is possible that the natural estrogen associations with functional connectivity may have been offset by estrogen therapy. Alternatively, this circuitry may be more sensitive to hormones in females rather than males, though no associations between hormones and behavior were seen. Previously, we have seen lower functional connectivity within the cerebellum associated with progesterone in females (Magalhães et al., 2025), which is also replicated here. It is possible that the CBBG circuit is more sensitive to progesterone due to hormonal transitions (menopause) occurring for the female participants in this age range, especially since the basal ganglia and cerebellum are susceptible to progesterone influences (Brinton et al., 2008; Pletzer et al., 2019). Because hormonal changes are robust and variable across this transition, brain circuits may alter their connectivity in response to these influences and show more sensitive responses to progesterone differences, especially as it is considered a neuroprotective hormone (Singh & Su, 2013). Further, as within-cerebellar connectivity differs with progesterone levels, the interactions between the cerebellum and basal ganglia may strengthen to offset these changes. Males do not experience the same progesterone fluctuations, so their circuits may be less sensitive to the hormone. The sex-dependent pattern of connectivity is interesting as some developmental DTI and functional connectivity studies have shown that males have greater within-hemisphere connectivity and females have greater between-hemisphere connectivity (Ingalhalikar et al., 2014). While the CBBG circuit is cross-hemispheric, here we are looking at patterns of connectivity between and within systems, rather than hemispheres.

A limitation of this study is the fact this it is cross-sectional, rather than longitudinal, which prevents us from interpreting any differences between age groups as causal. Longitudinal analyses are needed to better understand change in these circuits over time and with hormonal fluctuations. Another limitation is the inclusion of women in varying stages of menopause, including pre, peri, and post menopause. Women who are currently undergoing the menopausal transition have highly time variable hormones (Burger et al., 2002; Hale & Burger, 2009; Sherman & Korenman, 1975) which may not have been fully captured by one hormonal sample as was the case here.

## Conclusion

Overall, we found that the CBBG circuit can be parsed into subcircuits consistent with cortical networks and contributions of the regions, has differences in FC with increasing age and between sexes, and is associated with cognitive performance and progesterone levels. Though we did not find robust correlations between CBBG subcircuits and respective behavior composites, this circuitry remains a novel target to study in terms of cognitive and motor decline in healthy and pathological aging. These subcircuits can also be key areas to evaluate in cognitive and motor disorders, such as Parkinson’s and Alzheimer’s, to provide further insight into how brain connectivity is related to aging disorders that impact behavior. Though we did not find any significant connectivity between our cognitive and motor composites and their associated subcircuits, it is possible that these associations may be more prevalent in pathological aging, rather than healthy aging.

## Declarations

### Author Information

JAB designed the study. TH, TBA, and TM collected the data. TH and IH processed the data. CD and IH analyzed and interpreted the data and wrote the main manuscript text. JAB reviewed the manuscript.

## Acknowledgements

This work was supported by R01AG065010 to J.A.B. This work was further supported by the Texas Virtual Data Library (ViDaL), a high-performance cluster, funded by the Texas A&M University Research Development Fund.

## Data Availability Statement

The data that support the findings of this study are available from the corresponding author upon reasonable request.

## Ethical Approval

This study was performed in line with the principles of the Declaration of Helsinki. Approval was granted by the Texas A&M University Institutional Review Board.

## Consent To Participate

Informed consent was obtained from all individual participants included in the study.

