## Supplement for "Cerebello-basal ganglia networks and behavior across adulthood and aging"

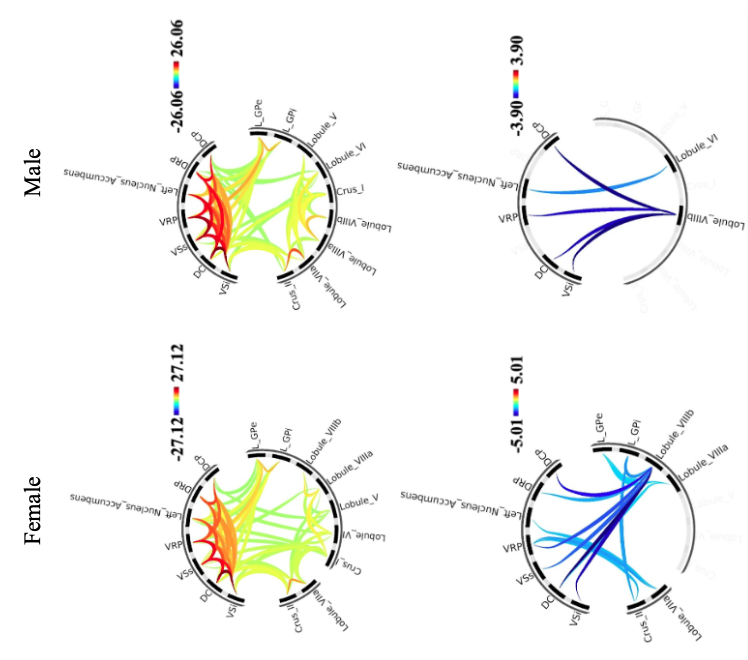

**Figure 1.** Patterns of functional connectivity (FC) of the CBBG circuit for each individual sex. The color bar represents the strength of the connectivity with red being positive and blue being negative. Results were evaluated using an analysis level FDR correction such that P < 0.05.

**Supplementary Table 1**. Functional connectivity (FC) between ROIs associated with cognitive behavior and networks.

| **FC Seed Regions** | **T(137)** | **pFDR** |
| --- | --- | --- |
| DC – VSi | 37.19 | 0.0000 |
| VRP – VSs | 31.59 | 0.0000 |
| LeftNAcc - VSs | 24.45 | 0.0000 |
| LeftNAcc - VRP | 23.37 | 0.0000 |
| VSs – DC | 28.31 | 0.0000 |
| VSs – VSi | 25.83 | 0.0000 |
| VRP – DC | 22.68 | 0.0000 |
| VRP – VSi | 19.60 | 0.0000 |
| LeftNAcc – DC | 19.46 | 0.0000 |
| LeftNAcc - VSi | 15.47 | 0.0000 |
| Crus II – DC | 8.18 | 0.0000 |
| Crus II – VSi | 7.75 | 0.0000 |
| Crus I – DC | 6.79 | 0.0000 |
| Crus I – VSi | 4.07 | 0.0000 |
| Crus II – Crus 1 | 6.40 | 0.0000 |
| Crus II – VSs | 4.67 | 0.0000 |
| Crus I – VSs | 4.47 | 0.0000 |
| Crus I – VRP | 2.10 | 0.0187 |
| Crus I - LeftNAcc | 2.80 | 0.0029 |
| Crus II – VRP | -1.85 | 0.0331 |

**Supplementary Table 2.** Functional connectivity (FC) between ROIs associated with motor behavior and networks.

| **FC Seed Regions** | **T(137)** | **pFDR** |
| --- | --- | --- |
| DRP - DCP | 24.12 | 0.0000 |
| LGPe – LGPi | 13.89 | 0.0000 |
| Lobule V – Lobule VI | 6.44 | 0.0000 |
| DRP – LGPe | 4.47 | 0.0000 |
| DCP – LGPe | 3.20 | 0.0009 |
| DCP - LGPi | 2.10 | 0.0188 |
| DRP – Lobule VI | 3.26 | 0.0008 |
| DCP – Lobule VI | 2.47 | 0.0073 |
| DCP – Lobule V | 1.80 | 0.0370 |

**Supplementary Table 3.** Functional connectivity (FC) of the cognitive subcircuit for each individual age group, including early middle age (EMA), late middle age (LMA), and older adulthood (OA).

| **FC Seed Regions for EMA** | **T(46)** | **pFDR** |
| --- | --- | --- |
| DC - VSi | 20.90 | 0.0000 |
| LeftNAcc - VSs | 19.56 | 0.0000 |
| LeftNAcc - VRP | 15.62 | 0.0000 |
| VRP - VSs | 18.47 | 0.0000 |
| VSs – DC | 18.55 | 0.0000 |
| VRP – DC | 14.64 | 0.0000 |
| VSs – VSi | 14.45 | 0.0000 |
| VRP - VSi | 11.91 | 0.0000 |
| LeftNAcc - DC | 11.63 | 0.0000 |
| LeftNAcc - VSi | 7.46 | 0.0000 |
| Crus II – DC | 4.54 | 0.0000 |
| Crus II – VSi | 4.07 | 0.0001 |
| Crus I – DC | 3.79 | 0.0002 |
| Crus I - VSi | 2.64 | 0.0057 |
| Crus II – Crus 1 | 4.66 | 0.0000 |
| Crus I – VSs | 4.36 | 0.0000 |
| Crus II – VSs | 2.77 | 0.0040 |
| Crus I – VRP | 1.79 | 0.0398 |
| Crus I - LeftNAcc | 3.76 | 0.0002 |
| Crus II - LeftNAcc | -1.79 | 0.0404 |
| **FC Seed Regions for LMA** | **T(50)** | **pFDR** |
| DC - VSi | 23.74 | 0.0000 |
| VSs – VSi | 19.74 | 0.0000 |
| VRP – VSs | 18.43 | 0.0000 |
| VRP - DC | 17.73 | 0.0000 |
| LeftNAcc – VSs | 17.46 | 0.0000 |
| VSs – DC | 16.75 | 0.0000 |
| LeftNAcc – VRP | 16.41 | 0.0000 |
| LeftNAcc - DC | 11.92 | 0.0000 |
| VRP - VSi | 11.73 | 0.0000 |
| LeftNAcc - VSi | 10.75 | 0.0000 |
| Crus II – DC | 6.02 | 0.0000 |
| Crus II - VSi | 5.29 | 0.0000 |
| Crus II - VSs | 4.63 | 0.0000 |
| Crus I - DC | 4.21 | 0.0001 |
| Crus I – VSi | 2.38 | 0.0214 |
| Crus I - VSs | 2.23 | 0.0306 |
| Crus I – Crus II | 3.34 | 0.0016 |
| **FC Seed Regions for OA** | **T(38)** | **pFDR** |
| VSi – DC | 19.81 | 0.0000 |
| VRP – VSs | 17.84 | 0.0000 |
| VSs - DC | 13.70 | 0.0000 |
| VSs – VSi | 11.37 | 0.0000 |
| VRP – VSi | 10.29 | 0.0000 |
| LeftNAcc – DC | 9.73 | 0.0000 |
| LeftNAcc – VRP | 9.58 | 0.0000 |
| LeftNAcc – VSs | 9.06 | 0.0000 |
| LeftNAcc – VSi | 8.98 | 0.0000 |
| VRP – DC | 8.79 | 0.0000 |
| Crus I – DC | 4.69 | 0.0000 |
| Crus II – VSi | 3.71 | 0.0003 |
| Crus II – DC | 3.33 | 0.0010 |
| Crus I - VSi | 2.09 | 0.0215 |
| Crus II – Crus I | 3.15 | 0.0016 |
| Crus II – VRP | -1.82 | 0.0383 |

**Supplementary Table 4.** Functional connectivity (FC) of the motor subcircuit for each individual age group, including early middle age (EMA), late middle age (LMA), and older adulthood (OA).

| **FC Seed Regions for EMA** | **T(46)** | **pFDR** |
| --- | --- | --- |
| DRP – DCP | 15.95 | 0.0000 |
| LGPi – LGPe | 9.66 | 0.0000 |
| Lobule VI – Lobule V | 3.31 | 0.0018 |
| DRP – Lobule VI | 2.57 | 0.0135 |
| DCP – Lobule VI | 2.13 | 0.03813 |
| DRP – LGPe | 2.34 | 0.0239 |
| DCP - LGPe | 2.23 | 0.0308 |
| **FC Seed Regions for LMA** | **T(50)** | **pFDR** |
| DRP – DCP | 15.71 | 0.0000 |
| LGPi – LGPe | 7.97 | 0.0000 |
| Lobule VI – Lobule V | 4.05 | 0.0002 |
| **FC Seed Regions for OA** | **T(38)** | **pFDR** |
| DRP – DCP | 10.94 | 0.0000 |
| LGPe – LGPi | 7.84 | 0.0000 |
| Lobule VI – Lobule V | 3.72 | 0.0007 |
| DRP – LGPe | 2.87 | 0.0067 |
| DRP - LGPi | 2.07 | 0.0454 |

**Supplementary Table 5.** Regions showing age differences in cerebello-basal ganglia functional connectivity.

| **FC Seed Regions** | **T(84)** | **pFDR** |
| --- | --- | --- |
| LGPe – VSs | 2.28 | 0.0126 |
| LGPe – DC | 1.93 | 0.0284 |
| LGPe – VSi | 1.86 | 0.0332 |
| LGPi - DC | 1.89 | 0.0308 |
| Crus I - LeftNAcc | 2.01 | 0.0238 |
| Lobule VIIIa – VSi | -3.18 | 0.0010 |
| Lobule VIIIa – VRP | -2.34 | 0.0109 |
| Lobule VIIIa - DRP | -1.79 | 0.0383 |
| Lobule VIIIa – DCP | -1.76 | 0.0414 |
| LGPi - VSs | -1.99 | 0.0247 |

**Supplementary Table 6.** Regions showing differences cerebello-basal ganglia functional connectivity associations with cognitive behavior across age

| **FC Seed Regions** | **T(134)** | **pFDR** |
| --- | --- | --- |
| Crus II – Lobule VI | 2.20 | 0.0146 |
| Lobule VIIIa – Crus I | -2.60 | 0.0052 |
| Crus I – Lobule VI | -2.16 | 0.0163 |
| Lobule VIIIa - LGPe | -1.71 | 0.0448 |
| Crus I – Lobule V | -1.67 | 0.0487 |

**Supplementary Table 7.** Regions showing sex differences in cerebello-basal ganglia functional connectivity.

| **FC Seed Regions** | **T(135)** | **pFDR** |
| --- | --- | --- |
| LeftNAcc – VRP | -3.40 | 0.0008 |
| LeftNAcc – VSi | -2.90 | 0.0044 |
| LeftNAcc – DC | -2.86 | 0.0049 |
| VRP – VSi | -2.95 | 0.0037 |
| VSs – VSi | -3.21 | 0.0037 |
| VRP – DC | -2.47 | 0.0146 |
| LeftNAcc – VSs | -2.37 | 0.0190 |
| DC – VSi | -2.33 | 0.0211 |
| VSs - DC | -2.46 | 0.0150 |
| DRP – LeftNAcc | -2.38 | 0.0189 |
| DRP – VRP | -2.24 | 0.0270 |
| DCP – VRP | -2.14 | 0.0342 |
| DCP - VSs | -2.14 | 0.0344 |

**Supplementary Table 8.** Regions showing Differences in cerebello-basal ganglia functional connectivity associations with progesterone across sex.

| **FC Seed Regions** | **T(134)** | **FC Seed Regions for EMA** |
| --- | --- | --- |
| LGPi – DRP | -1.76 | 0.0406 |
| LGPe – VSs | -1.68 | 0.0480 |
| Lobule VII – VRP | -1.71 | 0.0444 |
| LeftNAcc – VSi | -3.42 | 0.0004 |
| VRP – VSi | -3.05 | 0.0014 |
| DRP – VRP | -3.26 | 0.0007 |
| LeftNAcc – DC | -2.64 | 0.0046 |
| VRP – DC | -2.49 | 0.0069 |
| DRP – DC | -2.73 | 0.0036 |
| LeftNAcc – VRP | -2.36 | 0.0099 |
| DCP – VRP | -2.57 | 0.0056 |
| DCP – DRP | -2.37 | 0.0097 |
| DCP – VSs | -2.34 | 0.0104 |
| VRP – VSs | -1.85 | 0.0329 |
| VSs – DC | -2.20 | 0.0148 |
| VSs – VSi | -2.00 | 0.0240 |
| LeftNAcc – VSs | -1.74 | 0.0423 |
| Crus II – Lobule VIIIa | -2.88 | 0.0023 |
| Lobule VIIIb – Lobule V | -2.62 | 0.0050 |
| Crus II – LGPi | -1.94 | 0.0272 |
| Crus II – Lobule VI | -1.85 | 0.0331 |
| Crus II – Lobule VIIIb | -1.84 | 0.0342 |
| Lobule VIIIa – Crus I | -1.90 | 0.0300 |
| Lobule VIIIa – Lobule V | -1.90 | 0.0301 |
| Lobule VIIIb – Crus I | -1.98 | 0.0250 |
| Lobule VIIIa – Lobule VIIIb | -1.72 | 0.0435 |
| Lobule VIIIa – VRP | 3.69 | 0.0002 |
| Lobule VIIa – DRP | 3.16 | 0.0010 |
| Lobule VIIIa – VSs | 2.29 | 0.0117 |
| Crus I – VSs | 2.41 | 0.0086 |
| Crus I – LeftNAcc | 2.27 | 0.0124 |
| Crus I – DCP | 2.11 | 0.0183 |
| Crus I – VRP | 2.00 | 0.0238 |
| Lobule VIIIa – VSi | 1.71 | 0.0449 |
| Lobule VI – DC | 2.23 | 0.0136 |
| Lobule VI – VSs | 1.74 | 0.0423 |
| Lobule V - DRP | 1.99 | 0.0242 |
